# Lipid monounsaturation confers cold tolerance and improves tissue viability during hypothermic storage

**DOI:** 10.64898/2026.02.09.703646

**Authors:** Ran Zhang, Wenxuan Hou, Yuling Chen, Tong Nie, Yuxiang Tang, Zhiqiang Li, Changqing Zheng, Yupei Jiao, Xiaohui Liu, Ying Li, Jianlin Lei, Zhang Liu, Jianjun Wang, Qinghua Tao, Haiteng Deng

## Abstract

The rapid loss of tissue viability during hypothermic storage frequently results in organ failure and high discard rates in transplantation. In contrast, ectothermic animals can tolerate prolonged exposure to low temperatures in their natural environments. Elucidating the mechanisms underlying this cold tolerance may therefore inform improved strategies for tissue preservation. Here, we investigate proteomic and metabolomic responses to cold exposure in the livers of two frog species from distinct habitats: the cold-sensitive African clawed frog (*Xenopus laevis*) and the cold-tolerant Northeastern Asian brown frog (*Rana dybowskii*). Cold exposure induced lipid mobilization in *X. laevis*, whereas it promoted phospholipid mono-unsaturation in *R. dybowskii*. Notably, treatment with monounsaturated fatty acids (MUFAs) or overexpression of stearoyl-CoA desaturase (SCD) reduced cell death during cold storage. Moreover, MUFA infusion significantly improved cell viability in mouse liver tissue under hypothermic conditions. Together, these findings suggest that lipid mono-unsaturation, an adaptive feature of cold-tolerant frogs, can be leveraged to enhance tissue preservation during cold storage.

## INTRODUCTION

Organ and tissue transplantation saves millions of lives each year (Giwa et al., 2017). However, fewer than 10% of patients worldwide who require transplants ultimately receive them, largely due to the limited availability of donor organs (Ward et al., 2018). A major contributing factor is the lack of effective long-term preservation methods, which leads to high discard rates of donated organs (Giwa et al., 2017). Cryopreservation of large tissues and organs, such as livers and kidneys, remains particularly challenging, as ice formation during freeze-thaw cycles causes severe structural damage and requires the use of high concentrations of cytotoxic cryoprotective agents (CPAs). Consequently, static cold storage (SCS) at 0-4 °C remains the clinical standard for the preserving large tissues and organs for transplantation. Although hypothermic storage minimizes thermal stress and avoids the need for toxic CPAs, making it suitable for large and fragile organs (Huang et al., 2021), tissues preserved under these conditions deteriorate rapidly and are highly susceptible to ischemic injury. These limitations underscore the urgent need for improved preservation strategies that extend tissue and organ viability.

In contrast to human cells and tissues, which are highly vulnerable to cold-induced damage, many organisms have evolved remarkable mechanisms to survive prolonged exposure to extreme cold. For example, the Alaska wood frog (*Rana sylvatica*) can overwinter with much of its body frozen and fully recover upon thawing in the spring (Storey and Storey, 2017). Elucidating the protective mechanisms employed by such cold-tolerant organisms to maintain cellular and organ integrity at low temperatures may therefore inform new approaches to tissue and organ preservation (Liu et al., 2022). Endothermic animals, including humans, maintain stable body temperatures in cold environments through increased metabolic activity, whereas ectothermic animals, such as frogs, adapt to cold exposure by suppressing metabolic rates, enabling long-term survival under hypothermic or freezing conditions (Sokolova, 2019). This survival depends on extensive biochemical remodeling that preserves physiological function and prevents tissue damage. Studying these metabolic adaptations may thus reveal fundamental mechanisms of cold tolerance and inspire innovative preservation technologies.

The Northeastern Asian brown frog (*Rana dybowskii*) spends nearly half of the year in hibernation and exhibits exceptional cold tolerance. During hibernation, its metabolic rate, body temperature, and heart rate decline markedly, conserving energy and slowing biochemical processes (Zhang et al., 2020). Exposure to low temperatures also induces glycogen breakdown and elevates blood glucose levels, a response that may enhance freeze tolerance (Li et al., 2012; Xu et al., 2020). In contrast, the African clawed frog (*Xenopus laevis*), a widely used model organism in developmental biology, originates from the warm climates of sub-Saharan Africa and lacks comparable cold tolerance **(Figure 1A**). Systematic comparative analyses of cold-induced metabolic remodeling in these two species therefore provide a unique opportunity to uncover adaptive strategies underlying extreme cold tolerance and to inform the development of novel organ preservation approaches.

**Figure 1.**
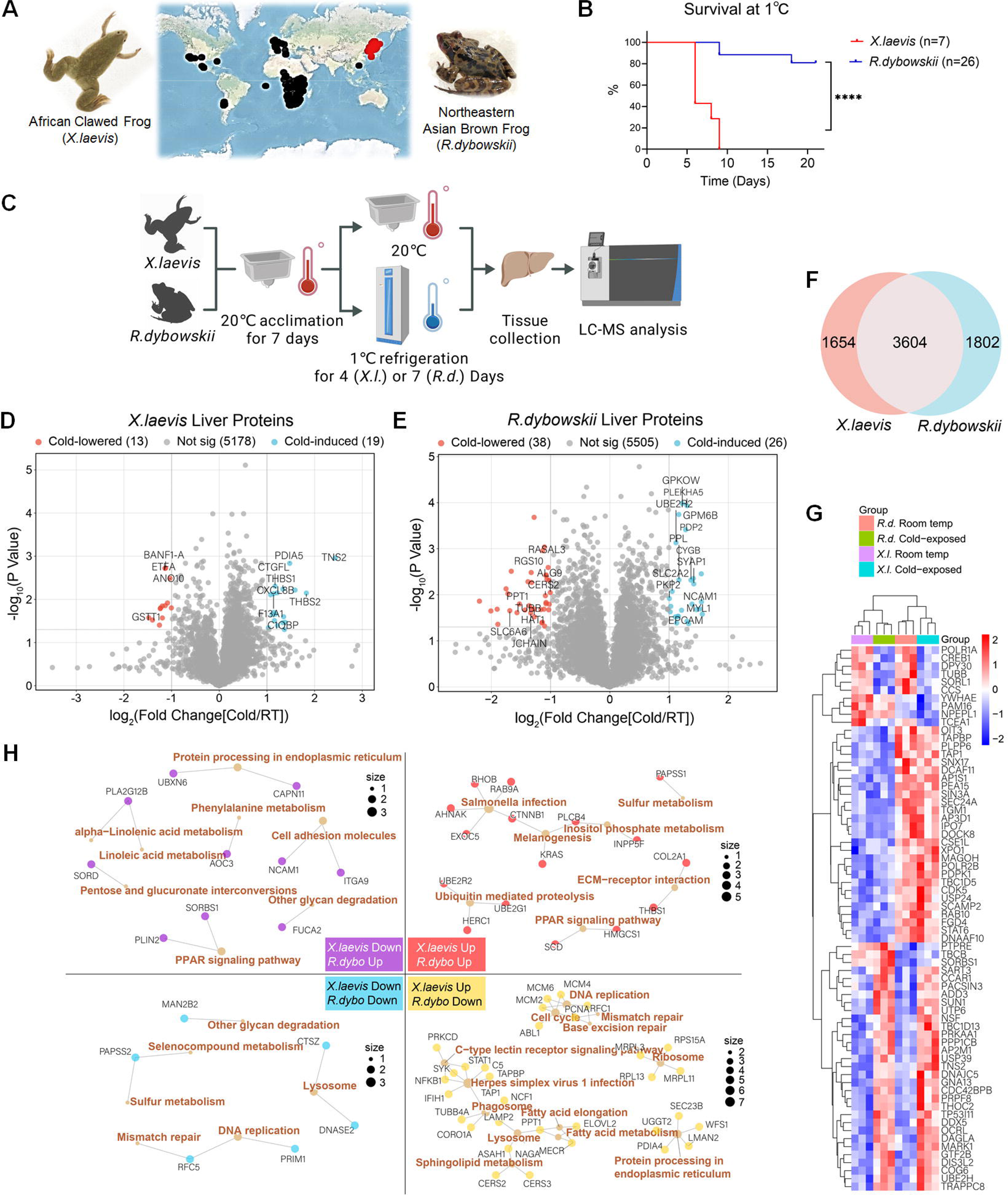
African clawed frog (*X. laevis*) and Northeastern Asian brown frog (*R. dybowskii*) respond differently to cold exposure. (A) The distribution area of the African clawed frog (*X. laevis*, on the left side of the map) is marked with black dots, while the distribution area of the Northeast Asian brown frog (*R. dybowskii*, on the right side of the map) is marked with red dots. Geographic distribution information is sourced from www.berkeleymapper.berkeley.edu. (B) Survival curves for African clawed frog and Northeast Asian brown frog maintained at 1 °C. ****p value < 0.0001 using Kaplan-Meier survival analysis. (C) Experimental workflow for proteomic and metabolomics analyses of two frog species upon cold exposure. (D) Volcano plot showing significantly upregulated (blue) and downregulated (red) proteins in *X. laevis* liver after cold exposure. (E) Volcano plot showing significantly upregulated (blue) and downregulated (red) proteins in *R. dybowskii* liver after cold exposure. (F) Venn diagram showing proteins identified separately and commonly in the liver proteomes of *X. laevis* and *R. dybowskii*. (G) Heatmap showing expression levels of differentially expressed proteins across species and environmental conditions, with hierarchical clustering and correlation analysis. (H) KEGG pathway network analysis generated using the igraph R package. Quadrant I (red): proteins and pathways upregulated in both species. Quadrant II (purple): proteins and pathways downregulated in *X. laevis* but upregulated in *R. dybowskii*. Quadrant III (blue): proteins and pathways downregulated in both species. Quadrant IV (yellow): proteins and pathways upregulated in *X. laevis* but downregulated in *R. dybowskii*. Protein names are shown in black; biological pathways are in bold brown. Layout is based on multidimensional scaling, with distances inversely proportional to pairwise Pearson correlation coefficients. Node size represents pathway enrichment magnitude.

In this study, we investigated cold-induced metabolic remodeling in the livers of both frog species using quantitative proteomic and metabolomic analyses. We further performed functional validation experiments in mammalian cells and mouse tissues to assess whether mimicking these metabolic adaptations enhances cell and tissue viability during low-temperature preservation. Together, our findings establish a framework for understanding species-specific cold adaptation strategies and identify supplementation with monounsaturated fatty acids as a promising approach for improving hypothermic tissue preservation.

## RESULTS

### African clawed frog (*X. laevis*) and Northeastern Asian Brown frog (*R. dybowskii*) respond differently to cold exposure

To assess cold tolerance, adult (∼2 years old) male African clawed frogs (*X. laevis*) and Northeastern Asian brown frogs (*R. dybowskii*) were acclimated at 20 °C for one week and then transferred to a refrigerator maintained at 1 °C. Survival was monitored daily. Under cold conditions, frogs of both species became immobile in the absence of external stimulation. However, all *X. laevis* individuals died within 6-9 days, whereas more than 80% of *R. dybowskii* remained alive and appeared largely healthy after 3 weeks at 1 °C, demonstrating markedly superior cold tolerance (**Figure 1B**).

To investigate molecular responses to cold exposure, we performed quantitative proteomic analyses of liver tissues following refrigeration at 1 °C for 4 days (*X. laevis*) or 7 days (*R. dybowskii*) (**Figure 1C**). In *X. laevis* livers, 7,863 protein groups were identified, of which 7,513 were quantified and 5,210 retained after filtering. Principal component analysis (PCA) showed tight clustering of cold-exposed samples, with limited separation from room-temperature controls (**Figure S1A**). Differential expression analysis identified 32 differentially expressed proteins (DEPs; |log₂FC| ≥ 1, p < 0.05), including 19 upregulated and 13 downregulated proteins (**Figure 1D**). Gene Ontology (GO) enrichment revealed that upregulated proteins were associated with testosterone response, cell spreading, and post-transcriptional regulation, whereas downregulated proteins were enriched in small-molecule, carbohydrate, and organic acid catabolic processes (**Figures S1C–D**). These results indicate relatively modest metabolic remodeling in *X. laevis* liver under cold exposure, primarily affecting lipid and carbohydrate metabolism.

In contrast, proteomic analysis of *R. dybowskii* livers identified 7,064 protein groups, with 6,910 quantified and 5,569 retained after filtering. PCA revealed clear separation between room-temperature and cold-exposed samples (**Figure S1B**). Differential expression analysis identified 64 DEPs, approximately twice the number observed in *X. laevis*, including 38 upregulated and 26 downregulated proteins (**Figure 1E**). Upregulated proteins were enriched in pathways related to actin cytoskeleton organization and cell migration, whereas downregulated proteins were associated with sphingolipid catabolism, membrane lipid metabolism, and neutrophil activation (**Figure S1E–F**). These findings suggest that *R. dybowskii* adapts to cold exposure through extensive cytoskeletal remodeling, lipid metabolic reprogramming, and suppression of innate immune activity.

To directly compare responses between species, protein identifiers were standardized, yielding 5,258 gene symbols for *X. laevis* and 5,406 for *R. dybowskii*, with 3,604 shared proteins (**Figure 1F**). Among these shared proteins, 68 were differentially expressed: 28 were downregulated and 6 upregulated in both species, while 34 exhibited opposite regulation patterns, indicating both conserved and species-specific cold responses (**Figure 1G**). After removing redundancies, 3,257 unique proteins were analyzed, of which 439 showed a fold change >1.5 (|log₂FC| > 0.58) in at least one species. KEGG pathway analysis revealed shared upregulation of pathways including ubiquitin-mediated proteolysis, extracellular matrix-receptor interaction, inositol phosphate metabolism, and PPAR signaling, while DNA replication and lysosomal pathways were downregulated in both species (**Figure 1H**). Notably, several pathways displayed opposing regulation: ER protein processing, cell adhesion, and PPAR signaling were upregulated in *R. dybowskii* but downregulated in *X. laevis*, whereas pathways associated with DNA replication, cell cycle progression, ribosome function, ER protein processing, and immune signaling were upregulated in *X. laevis* but suppressed in *R. dybowskii* (**Figure 1H**). Together, these data indicate fundamentally different cold-adaptation strategies. *X. laevis* responds to acute cold stress by activating energy-intensive processes, including cell proliferation, protein synthesis, and immune signaling, whereas *R. dybowskii* conserves energy during prolonged cold exposure by suppressing high-demand pathways and remodeling cytoskeletal and lipid metabolic programs.

### Metabolomic analysis of African clawed frog and Northeastern Asian Brown frog liver upon cold exposure

To characterize metabolic remodeling in response to cold exposure, we performed untargeted metabolomic profiling of liver tissues from African clawed frogs (*X. laevis*) and Northeastern Asian brown frogs (*R. dybowskii*). In *X. laevis*, 573 metabolites were identified in positive ion mode and 340 in negative ion mode, with 74 metabolites detected in both modes (**Figure 2A**). PCA showed limited separation between room-temperature (RT) and cold-exposed (CE) samples, indicating minimal global metabolic changes in *X. laevis* liver following cold exposure (**Figure 2B**). In contrast, metabolomic profiling of *R. dybowskii* identified 662 metabolites in positive mode and 358 in negative mode, with 84 shared between modes (**Figure 2A**). PCA revealed clear separation between RT and CE samples, consistent with a robust metabolic response to cold exposure in *R. dybowskii* liver (**Figure 2B**). Differential metabolite analysis in *X. laevis* identified 44 significantly upregulated and 4 downregulated metabolites (|log₂FC| ≥ 1, p < 0.05) (**Figure 2C**). Pathway enrichment analysis indicated involvement of arginine and proline metabolism, pyruvate metabolism, linoleic acid metabolism, and the tricarboxylic acid (TCA) cycle (**Figure 2D**). Notably, several free fatty acids (e.g., FA(20:2), FA(26:5)) and acylcarnitines, including palmitoylcarnitine, oleoylcarnitine, and arachidonylcarnitine, were significantly upregulated, suggesting enhanced lipid mobilization and fatty acid oxidation to meet energy demands during cold exposure (**Figure 2C**).

**Figure 2.**
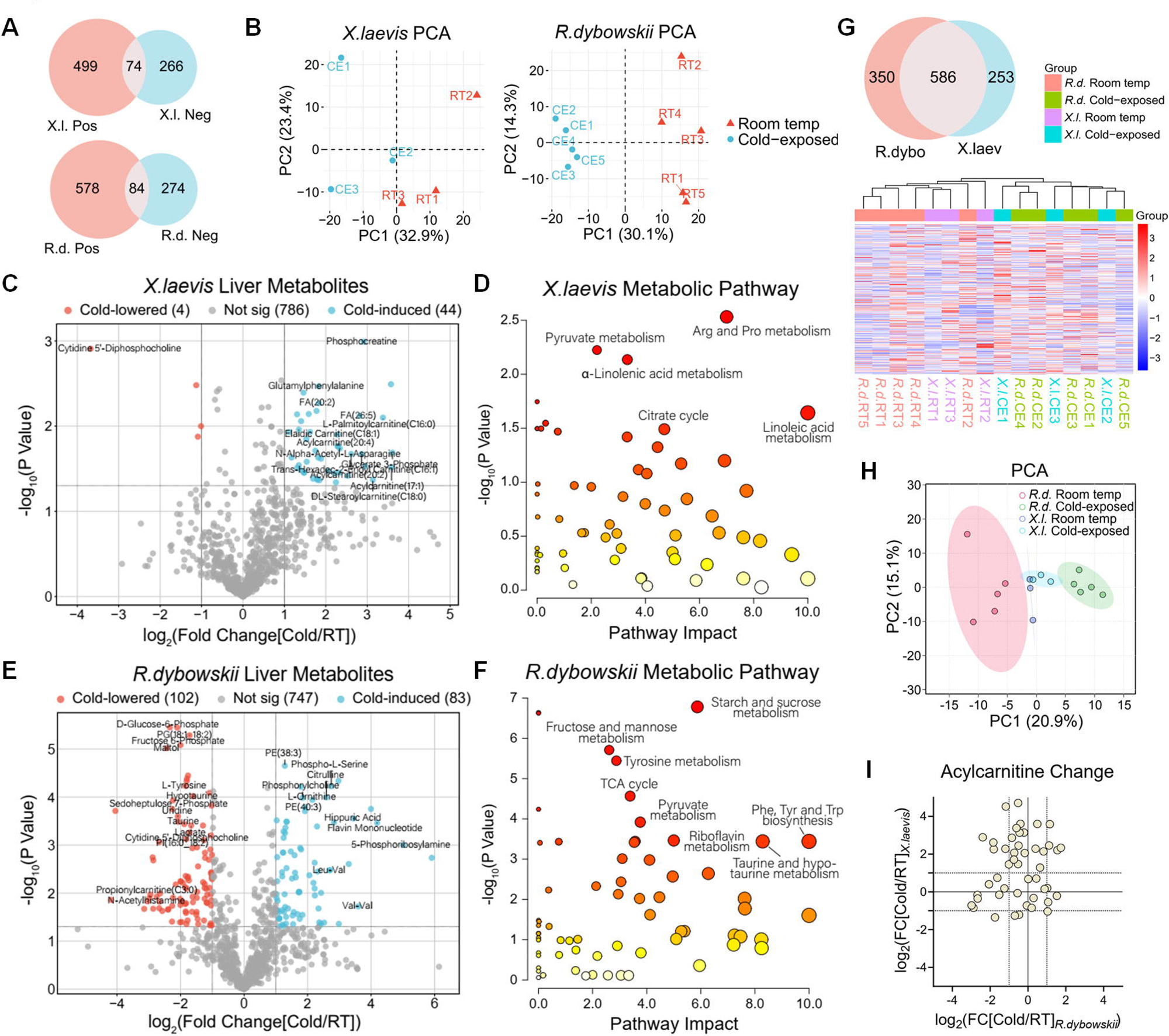
Metabolomic analysis of African clawed frog and Northeastern Asian brown frog liver upon cold exposure. (A) Venn diagram showing compounds quantified in positive and negative ion modes, separately and commonly, in African clawed frog livers (upper) and Northeastern Asian brown frog livers (lower), respectively. (B) PCA of African clawed frog livers (left) and Northeastern Asian brown frog livers (right) showing the relative distances between samples. (C and E) Volcano plot showing significantly upregulated (blue) and downregulated (red) metabolites in African clawed frog livers (C) and Northeastern Asian brown frog livers (E), respectively, after cold exposure. (D and F) Metabolic pathway enrichment analysis of African clawed frog livers (D) and Northeastern Asian brown frog livers (F) using MetaboAnalyst. (G) Venn diagram showing compounds identified separately and commonly in both species’ liver metabolomes. Heatmap showing expression levels of commonly identified compounds across species and environmental conditions, with hierarchical clustering and correlation analysis. (H) PCA showing relative distances between samples. (I) Log_2_ fold changes of acylcarnitines after cold exposure in African clawed frog (*X. laevis*, y-axis) and Northeastern Asian brown frog (*R. dybowskii*, x-axis) liver. Each point represents one acylcarnitine.

In *R. dybowskii*, 83 metabolites were significantly upregulated and 102 downregulated (|log₂FC| ≥ 1, p < 0.05) (**Figure 2E**). Enriched pathways included carbohydrate metabolism (starch and sucrose, fructose and mannose), amino acid metabolism (tyrosine and aromatic amino acids), energy metabolism (pyruvate metabolism and the TCA cycle), and cofactor metabolism (riboflavin, taurine, and hypotaurine) (**Figure 2F**). In addition, substantial phospholipid remodeling was observed: phosphatidylethanolamines (e.g., PE(38:3), PE(40:3)) and phosphatidylcholines were upregulated, whereas phosphatidylglycerols (PG(18:1_18:2)) and phosphatidylinositols (PI(16:0_18:2)) were downregulated, indicating altered membrane lipid composition under cold stress (**Figure 2E**).

Comparative analysis revealed that 586 metabolites were shared between the two species (**Figure 2G**). Hierarchical clustering showed that metabolic profiles clustered more closely by temperature than by species, with cold exposure driving similar shifts in both frogs (**Figures 2G–H**). However, separation between RT and CE samples was substantially greater in *R. dybowskii*, indicating more extensive metabolic remodeling (**Figure 2H**). Despite widespread upregulation of free fatty acids and acylcarnitines in X. laevis, these metabolites remained largely unchanged or were slightly downregulated in *R. dybowskii* (**Figures 2I** and **S2**), suggesting minimal lipid mobilization in the cold-adapted species.

Instead, integrated proteomic and metabolomic analyses revealed enhanced lipid monounsaturation in *R. dybowskii*. Protein expression of stearoyl-CoA desaturase (SCD), a key enzyme that converts saturated fatty acids (SFAs) into monounsaturated fatty acids (MUFAs) by introducing a cis double bond at the Δ9 position, was upregulated in cold-exposed livers (**Figure 3A**). Consistently, the MUFA/SFA ratio in phospholipids increased in *R. dybowskii* (**Figure 3B**). In contrast, *X. laevis* showed only modest SCD upregulation and no clear shift in MUFA/SFA ratios (**Figures S3A–B**), indicating that *R. dybowskii* selectively remodels membrane phospholipids to increase monounsaturation as a key cold-adaptive strategy.

**Figure 3.**
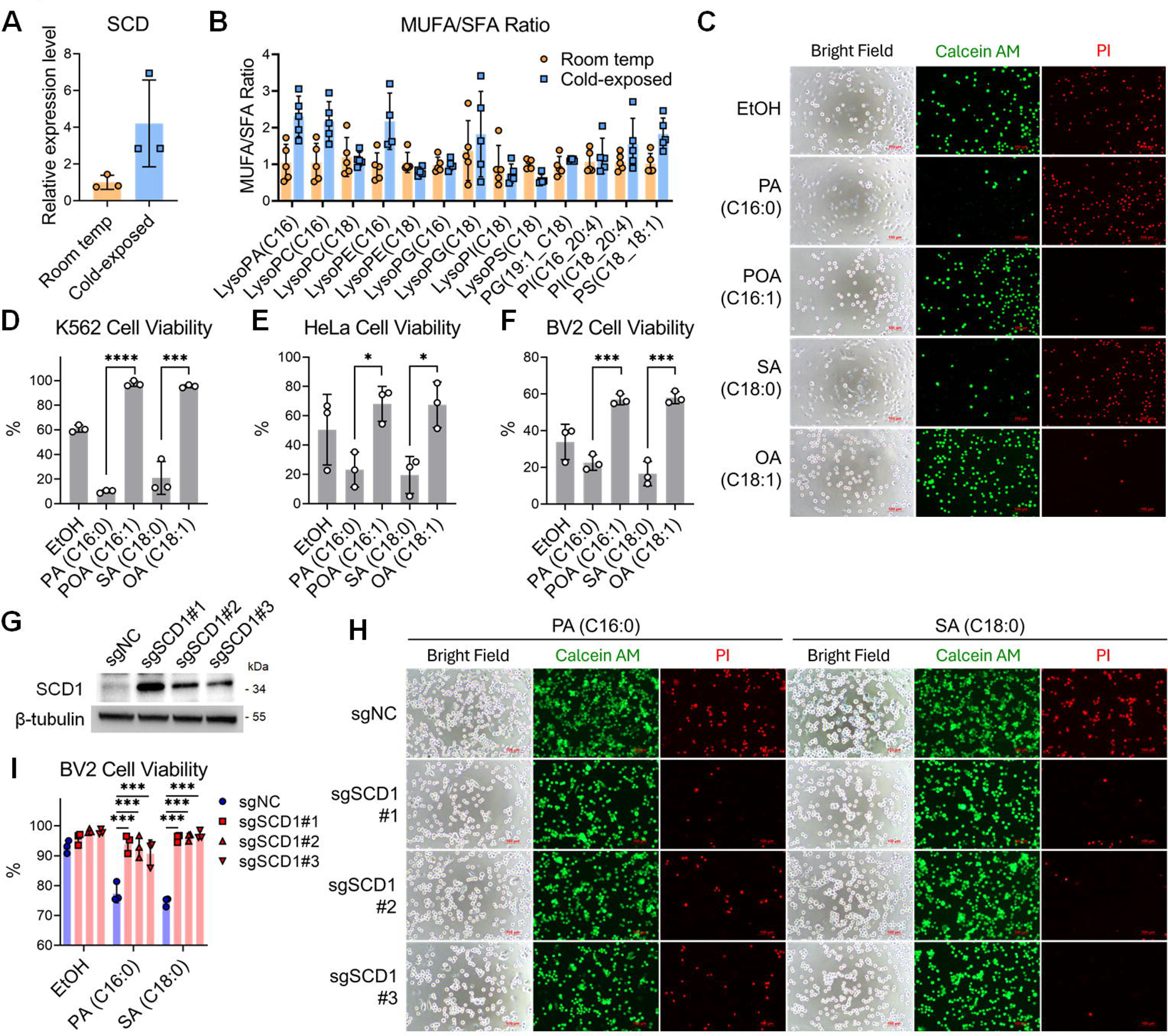
MUFA improves cell survival under cold conditions. (A) Relative SCD protein expression levels in Northeastern Asian brown frog liver measured by quantitative proteomic analysis. N = 3. (B) Relative ratios of monounsaturated to saturated phospholipids (MUFA/SFA) in Northeastern Asian brown frog quantified through metabolomic analysis. N = 3. (C) Calcein AM/PI double staining and representative microscopy imaging showing survival (green) and death (red) of K562 cells pretreated with different fatty acids followed by cold exposure. (D-F) Cell survival rate in K562 (D), HeLa (E), and BV2 (F) cells pretreated with different fatty acids followed by cold exposure, quantified. N = 3. (G) Western blot analysis of SCD1 protein expression levels in control (sgNC) and SCD1-overexpressing (sgSCD1#1-3) BV2 CRISPRa cells. (H and I) Calcein AM/PI staining showing survival (green) and death (red) of control and SCD1-overexpressing BV2 CRISPRa cells pretreated with PA or SA and exposed to cold (H). The quantification of the survival rates is shown in (I). N = 3. Data are presented as mean ± SD. Statistical significance was determined using unpaired Student’s t-test (D-F), and two-way ANOVA with Sidak’s multiple comparisons (I). *p < 0.05, ***p < 0.001, ****p < 0.0001.

### MUFA supplementation and SCD1 overexpression improve cell survival under cold stress

Given the critical role of phospholipid saturation in determining membrane biophysical properties and in mediating communication between intracellular and extracellular environments, we asked whether fatty acid monounsaturation influences cell survival under cold stress. To address this question, we evaluated the effects of long-chain fatty acids on cell viability at low temperature. Suspension (K562), adherent (HeLa), and semi-adherent (BV2) cells were pretreated with individual fatty acids for 6 h at 37 °C and then exposed to 4 °C overnight (∼16 h). Calcein AM/PI staining showed that SFAs, including palmitic acid (PA) and stearic acid (SA), increased cell death across all three cell types, whereas MUFAs, including palmitoleic acid (POA) and oleic acid (OA), significantly reduced cell death and improved survival under cold conditions (**Figures 3C-F** and **S4A-B**). Notably, MUFA-mediated protection required several hours of pretreatment, as immediate cold exposure after fatty acid addition did not confer protection, suggesting that MUFAs act through cellular metabolic processes, such as incorporating into membrane phospholipids, rather than exerting an immediate effect.

To directly test the role of endogenous fatty acid monounsaturation, we generated BV2 cell lines overexpressing stearoyl-CoA desaturase 1 (SCD1) using CRISPR activation. Western blotting confirmed robust upregulation of SCD1 protein in sgSCD1#1–3 cells compared with non-targeting controls (sgNC) (**Figure 3G**). Upon PA or SA pretreatment followed by cold exposure, SCD1-overexpressing cells exhibited significantly reduced cell death relative to control cells (**Figures 3H–I**), indicating that SCD1-mediated protection depends on the conversion of SFAs into MUFAs.

Because fatty acid saturation influences oxidative vulnerability, we next examined lipid peroxidation and oxidative stress under cold conditions. BODIPY 581/591 C11 staining showed that SFA treatment markedly increased lipid peroxidation after cold exposure, whereas MUFA treatment suppressed oxidation (**Figures 4A–B**). Consistently, DCFH-DA staining demonstrated that cold exposure elevated intracellular reactive oxygen species (ROS), which were further enhanced by SFAs but attenuated by MUFAs (**Figure 4B**). Measurements of malondialdehyde (MDA), a lipid peroxidation byproduct, corroborated these findings (**Figure 4C**). At the signaling level, cold exposure induced phosphorylation of p38, RIP3, and MLKL. SFA treatment further amplified RIP3 and MLKL phosphorylation, whereas MUFA treatment suppressed necroptotic signaling under cold conditions (**Figure 4D**). In addition, fluorescence polarization measurements using TMA-DPH revealed that MUFA treatment increased membrane fluidity, providing a potential biophysical basis for enhanced cold tolerance (**Figure 4E**).

**Figure 4.**
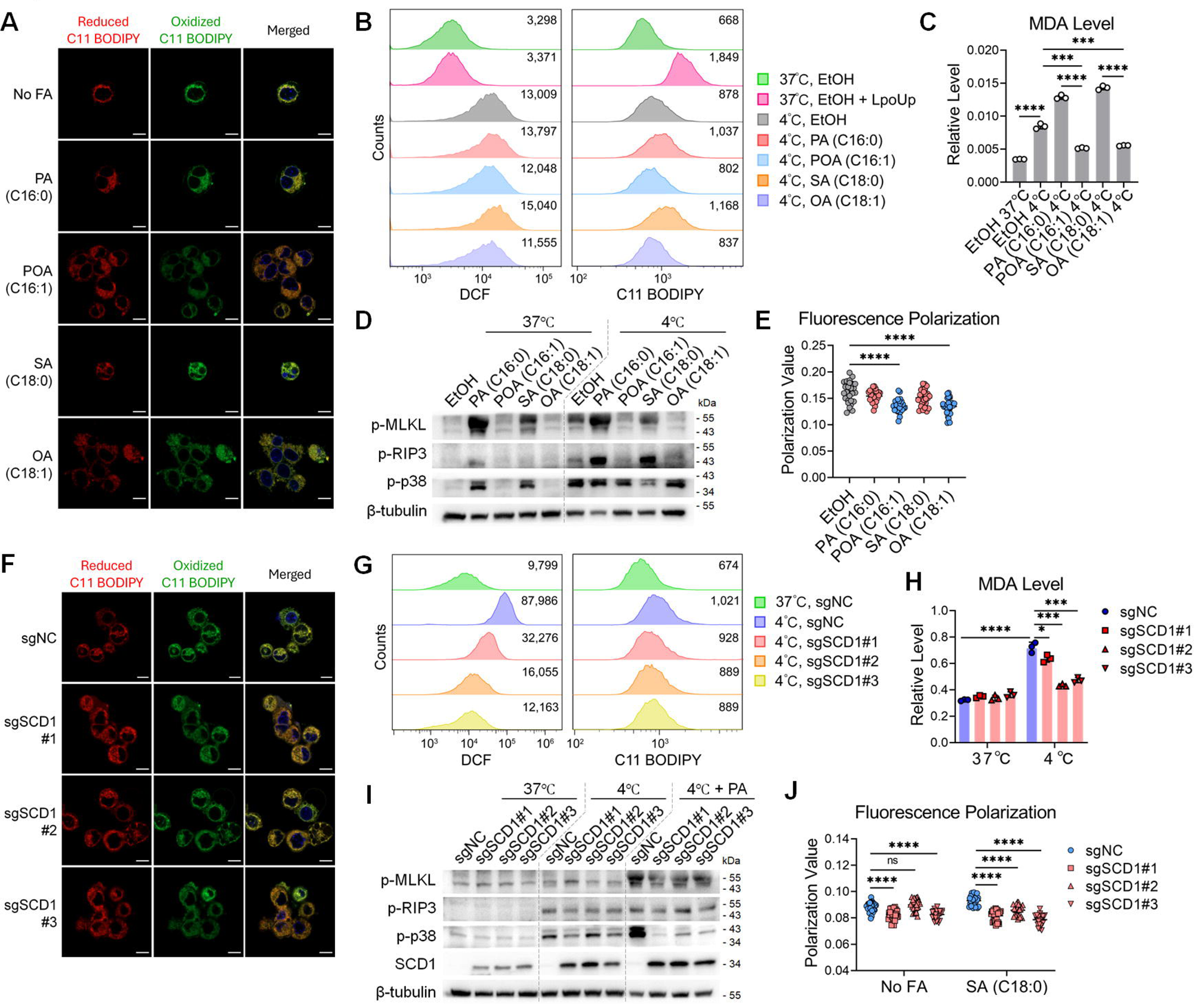
MUFA treatment reduces lipid peroxidation and improves membrane fluidity under cold conditions. (A) Confocal imaging of BV2 cells stained with BODIPY 581/591 C11 after fatty acid pretreatment and cold exposure. (B) Flow cytometry analysis of oxidative stress and lipid peroxidation using DCFH-DA (left column) or BODIPY 581/591 C11 (right column). Mean fluorescence values are indicated in the top right corner. (C) MDA levels in BV2 cells after fatty acid pretreatment and cold exposure. N = 3. (D) Western blot analysis of death signaling pathways in BV2 cells after fatty acid pretreatment. (E) TMA-DPH staining and fluorescence polarization analysis. N = 27. (F) Confocal imaging of BODIPY 581/591 C11 staining in control and SCD1-overexpressing BV2 cells after fatty acid pretreatment and cold exposure. (G) Flow cytometry of DCFH-DA (left column) or BODIPY 581/591 C11 (right column) staining. Mean fluorescence values are in the top right corner (H) MDA levels in control and SCD1-overexpressing BV2 cells. N = 3. (I) Western blot analysis of death signaling pathways in control and SCD1-overexpressing BV2 cells. (J) TMA-DPH staining and polarization analysis. N = 27. Data are presented as mean ± SD. Statistical significance was determined using unpaired Student’s t-test (C and E) and two-way ANOVA with Sidak’s test (H and J). NS: not significant, ***p < 0.001, ****p < 0.0001.

We next assessed whether SCD1 overexpression recapitulates the protective effects of MUFAs. Under cold exposure, SCD1-overexpressing cells displayed significantly lower levels of ROS and lipid peroxidation compared with control cells, as measured by DCFH-DA and oxidized BODIPY C11 fluorescence (**Figures 4F–G** and **S5**). MDA levels were similarly reduced in SCD1-overexpressing cells following cold exposure (**Figure 4H**). Western blot analysis showed that SCD1 overexpression attenuated cold- and SFA-induced phosphorylation of p38 and MLKL (**Figure 4I**). Moreover, TMA-DPH polarization assays indicated that SCD1 overexpression increased membrane fluidity independently of SFA supplementation (**Figure 4J**). Together, these results demonstrate that fatty acid monounsaturation, achieved either through MUFA supplementation or SCD1 overexpression, protects cells from cold-induced injury by reducing oxidative stress and lipid peroxidation, suppressing necroptotic signaling, and enhancing membrane fluidity.

### Tissue degradation during cold storage of mouse liver

To translate cold-adaptive mechanisms identified in the Northeastern Asian brown frog to mammalian organ preservation, we first characterized tissue degradation during cold storage of mouse liver. Livers from 5-week-old mice were harvested and stored in DMEM at 4 °C for varying durations. Histological analysis revealed progressive structural deterioration with increasing storage time: cytoplasmic components of hepatocytes gradually diminished, as indicated by reduced eosin staining, and the hepatic cord architecture became increasingly disorganized, consistent with progressive cellular degradation during cold preservation (**Figure 5A**). TUNEL staining further showed a marked increase in apoptotic cells after 48 h of cold storage (**Figures 5A–B**).

**Figure 5.**
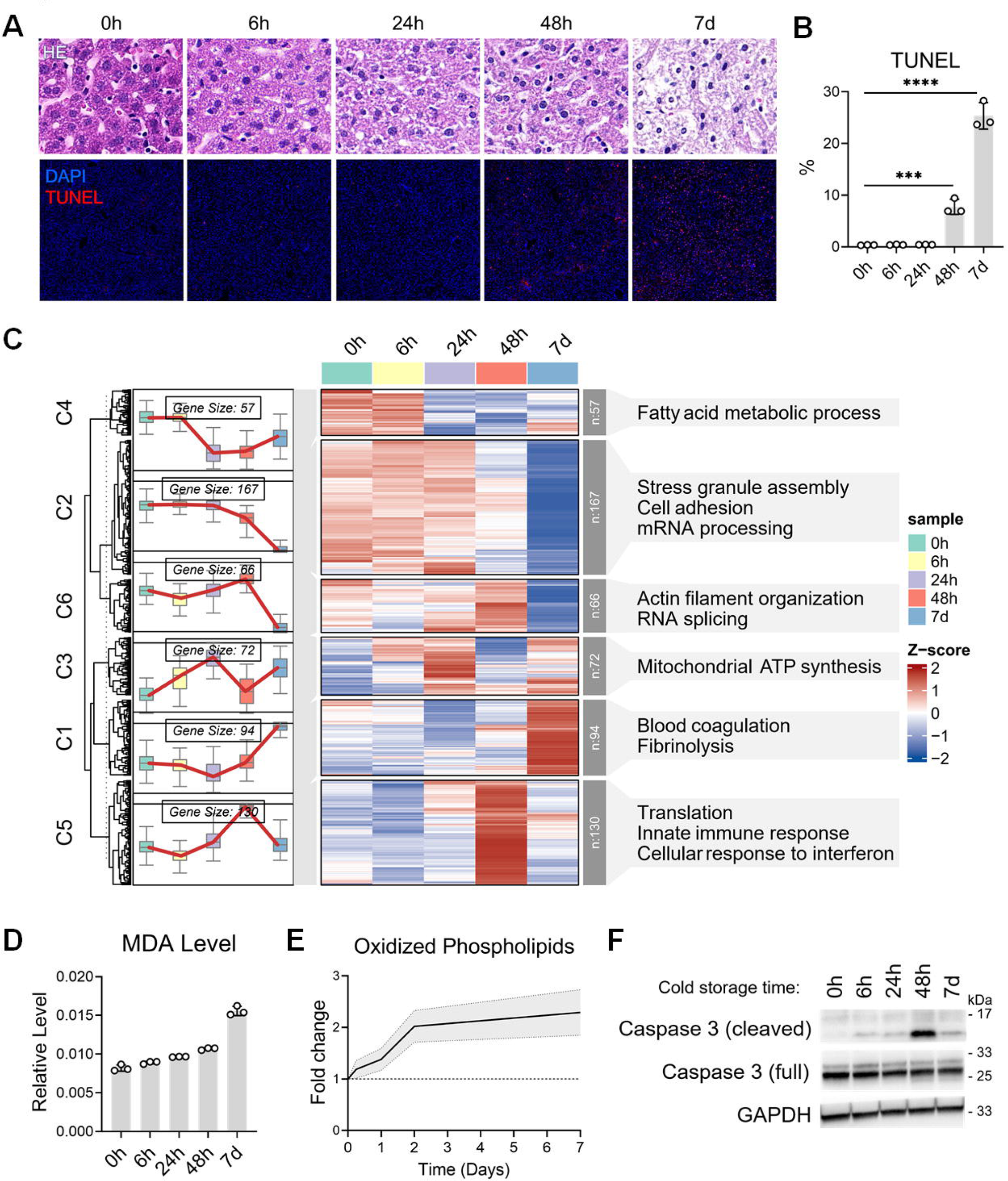
Tissue degradation in mouse liver during cold storage. (A) HE staining (top row) and TUNEL fluorescence microscopy (bottom row, with nuclei shown using DAPI staining) showing histopathology of mouse liver tissues at various time points after cold storage. (B) Quantification of the percentage of TUNEL-positive cells. N = 3. (C) Proteomic analysis of changes in mouse liver tissues during cold storage. Heatmap showing hierarchical clustering of proteins based on expression trends over time during cold storage, divided into six clusters (C1-6, leftmost). The number of differentially expressed genes in each cluster and the enriched biological processes through GO analysis are shown on the right side. (D) MDA levels in mouse liver tissues at various time points after cold storage. N = 3. (E) Relative levels of oxidized phospholipids normized to the non-oxidized forms were measured through lipidomic analysis. N = 9. (F) Western blot analysis of cleaved caspase 3 levels in mouse liver tissues at various times after cold storage. Data are presented as mean ± SD (B and D), or mean with 95% CI (E). Statistical significance was determined using unpaired Student’s t-test (B). ***p < 0.001, ****p < 0.0001.

To define molecular changes underlying tissue degradation, we performed quantitative proteomic analysis of mouse livers stored for different durations, identifying and quantifying 5,535 proteins. Temporal profiling revealed distinct stages of biological disruption during cold preservation. Within the first 24 h, proteins involved in mitochondrial ATP synthesis were upregulated (**Figure 5C**). By 48 h, proteins associated with essential cellular processes, including cell adhesion, mRNA processing, and stress granule assembly, were substantially downregulated (**Figure 5C**). Notably, proteins involved in translation, innate immune responses, and interferon signaling exhibited transient upregulation at 48 h, followed by a subsequent decline (**Figure 5C**). In contrast, proteins related to blood coagulation and fibrinolysis were markedly upregulated from 48 h onward, indicating extensive tissue and protein degradation during prolonged cold storage (**Figure 5C**).

Complementary lipidomic analyses revealed pronounced membrane lipid remodeling during cold preservation. Lysophospholipid levels increased progressively over time (**Figure S6A**). Phosphatidic acid (PA) levels increased within the first 48 h and then declined (**Figure S6B**), consistent with a sequential phospholipid degradation process in which head-group cleavage initially generates PA, followed by sn-2 acyl chain hydrolysis to produce lysophospholipids. Cardiolipin (CL) levels increased within the first 24 h and then decreased through day 7 (**Figure S6C**). In contrast, major plasma membrane phospholipids, including phosphatidylcholine (PC), phosphatidylethanolamine (PE), phosphatidylserine (PS), phosphatidylglycerol (PG), phosphatidylinositol (PI), and sphingomyelin (SM), underwent rapid depletion during the early storage phase (6–24 h), followed by a transient rebound at 48 h and subsequent stabilization or decline (**Figures S6D–I**). Given that CL is localized exclusively to mitochondrial membranes, these results indicate that plasma membrane phospholipids are more susceptible to degradation than mitochondrial membrane lipids during cold preservation, with the exception of the transient 48 h response.

We next investigated lipid peroxidation during cold storage. MDA levels steadily increased over time (**Figure 5D**), and consistently, lipidomics analysis showed a general rise in oxidized phospholipid species throughout the preservation period (**Figure 5E**). Moreover, Western blot analysis revealed increased levels of cleaved caspase-3 as early as 6 h of storage, peaking at 48 h and declining thereafter, likely due to extensive protein degradation at later time points (**Figure 5F**). Together, these data demonstrate that cold storage of mouse liver induces progressive phospholipid oxidation, membrane degradation, and apoptotic activation, ultimately compromising tissue integrity.

### Palmitoleic acid infusion reduces tissue damage in mouse liver during cold storage

To assess the effects of MUFA treatment on liver viability during cold storage, we injected 4-week-old male ICR mice (approximately 30 g body weight) with palmitoleic acid (POA; 250 µL of 5 mM solution) via the tail vein, 6 hours before liver harvest. After euthanasia, whole-body perfusion was performed using University of Wisconsin (UW) solution containing 250 µM POA. The liver was excised, immersed in 10 mL of pre-chilled UW solution supplemented with 250 µM POA, and stored at 4 °C for 48 hours.

Lipidomic analysis revealed that POA treatment decreased the percentages of lysophospholipids and CL, while increasing PC and PE levels, indicating reduced membrane degradation compared to controls (**Figure 6A**). Additionally, the POA-treated liver had a higher proportion of palmitoleic acid (C16:1) incorporated into PE (**Figure 6B**). To assess tissue damage, we measured hepatic transaminases released into the preservation solution. ALT levels were consistently lower in the POA-treated group than in controls at 0, 2, and 5 days of storage (**Figure 6C**), and AST levels were lower in the POA-treated group after 2 days (**Figure 6D**). Histological analysis showed less cellular degradation in POA-treated livers compared to controls (**Figure 6E**), with TUNEL staining revealing significantly fewer apoptotic cells in the POA-treated group (**Figure 6E**). Western blot analysis also indicated reduced levels of cleaved caspase-3 in POA-treated livers (**Figure 6F**). These results demonstrate that POA preconditioning, combined with POA supplementation during storage, reduces tissue degradation during liver cold preservation.

**Figure 6.**
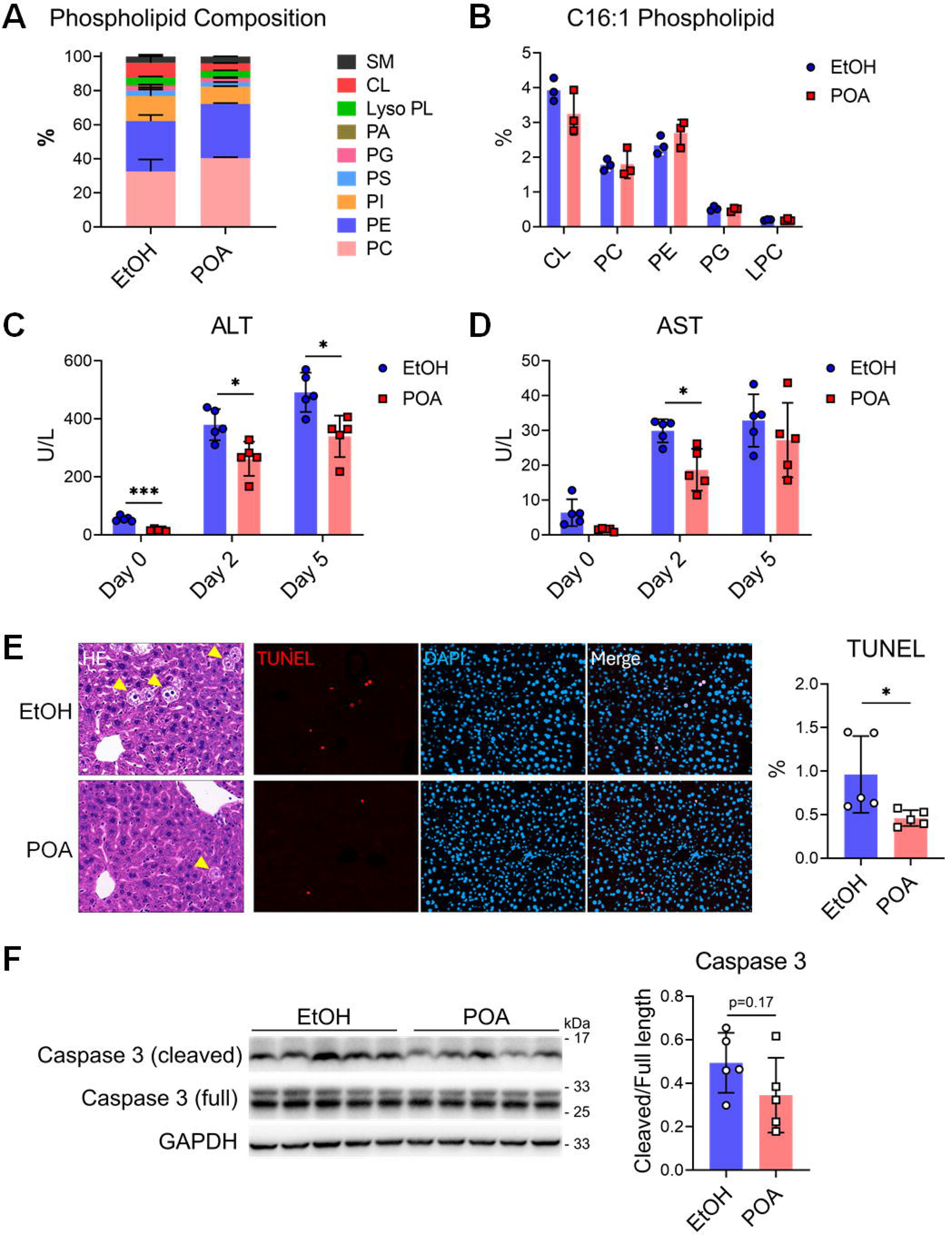
POA treatment reduces tissue damage in mouse liver during cold storage. (A) Phospholipid composition in mouse liver tissues after 48 h of cold storage with or without POA supplementation measured through lipidomic analysis. N = 3. (B) Percentage of palmitoleic (C16:1) acyl chain-containing phospholipids in each type of phospholipid measured through lipidomic analysis. N = 3. (C and D) ALT (C) and AST (D) levels released into the preservation solution at different time points during mouse liver cold storage. N = 5. (E) HE and TUNEL imaging of mouse liver tissues after 48 hours of cold storage treated with POA or ethanol vehicle. Nuclei were shown with DAPI staining. Yellow arrowheads indicate damaged cells. The percentages of TUNEL-positive cells were quantified. N = 5. (F) Western blot of cleaved caspase 3 in mouse liver tissues after 48 hours of cold storage with POA or vehicle treatment. The optical densities of cleaved caspase 3 relative to its full-length form were quantified. N = 5. Data are presented as mean ± SD. Statistical significance was determined using two-way ANOVA with Sidak’s test (C and D), and unpaired Student’s t-test (E and F). *p < 0.05.

In addition to the liver, we examined the effects of POA on other tissue types by cold-preserving excised dorsal skin from mouse ears in pre-chilled UW solution containing 250 µM POA at 4 °C for 48 hours. Compared with controls, POA-treated skin exhibited a thicker deep dermal layer and markedly reduced tissue degradation (**Figure S7**). Consistently, TUNEL staining revealed significantly fewer apoptotic cells in POA-treated samples (**Figure S7**). Together, these results suggest that POA has broad potential for mitigating tissue degradation during cold preservation.

## DISCUSSION

In this study, quantitative proteomic and metabolomic analyses revealed that two frog species from distinct natural habitats exhibit divergent responses to low-temperature stress. The relatively cold-sensitive African clawed frog displayed increased lipid mobilization and catabolism, elevated protein synthesis, and activation of innate immune pathways upon cold exposure, consistent with a strategy aimed at maintaining physiological homeostasis through increased energy expenditure. In contrast, the cold-tolerant Northeastern Asian brown frog exhibited suppression of protein synthesis and innate immune activity, reflecting an energy-conserving strategy that supports long-term survival under cold conditions. Elucidating the adaptive metabolic remodeling of the Northeastern Asian brown frog may therefore inform the development of more effective strategies for tissue and organ preservation.

Stearoyl-CoA desaturase (SCD) is the key enzyme that introduces a double bond at the Δ9 position of long-chain saturated fatty acids, thereby directly regulating the synthesis of MUFAs (Enoch et al., 1976). Integrated proteomic and metabolomic analyses revealed that, in the Northeastern Asian brown frog, SCD expression was upregulated after cold exposure, accompanied by an increase in the ratio of MUFA- to SFA-containing phospholipids (MUFA/SFA). In contrast, SCD upregulation in African clawed frog livers was modest, and was not associated with an increase in the MUFA/SFA phospholipid ratio. Interestingly, previous studies have shown that hepatic SCD1 expression in mice decreases significantly after 24 hours of cold exposure at 4 °C (Grefhorst et al., 2018). Together, these findings highlight species-specific regulation of hepatic SCD in response to cold stress: SCD expression decreases in endothermic animals (e.g., mice), increases in cold-tolerant ectothermic animals (e.g., Northeastern Asian brown frogs), and shows an intermediate response in cold-sensitive ectotherms (e.g., African clawed frogs).

We further examined the functional effects of fatty acids on cell survival by treating cells with individual fatty acids and generating SCD1-overexpressing cell lines. SFAs, such as PA and SA, exacerbated cell death under cold conditions, whereas MUFAs, including POA and OA, reduced cell death under the same conditions. Moreover, SCD1 overexpression mitigated cold-induced cell death in the presence of exogenous SFAs. Mechanistically, both MUFA treatment and SCD1 overexpression reduced ROS and lipid peroxidation levels in cells under cold stress, concomitantly attenuating the activation of stress- and cell death-associated signaling pathways. Consistent with these *in vitro* findings, *in vivo* experiments demonstrated that POA infusion and supplementation in preservation solution protected mouse liver and skin tissues from cold-induced cell death. Collectively, these results indicate that MUFAs enhance tissue and organ viability during low-temperature preservation.

Cold-tolerant frogs exhibit seasonal remodeling of membrane phospholipid composition to maintain cellular integrity under low-temperature conditions. Previous studies have shown that, compared with summer, overwintering wood frogs display increased hepatic PE levels and reduced PS or PC levels, changes that enhance membrane fluidity and facilitate cold adaptation (Reynolds et al., 2014). Consistent with these observations, our metabolomic analysis revealed a significant increase in hepatic PE levels in Northeastern Asian brown frogs following cold exposure, including elevations in PE(34:1), PE(p36:5), PE(38:3), PE(40:4), PE(40:5), and PE(40:3). Membrane fluidity is strongly influenced by fatty acid saturation: SFA chains are relatively linear and promote tight phospholipid packing, whereas unsaturated fatty acid chains introduce curvature and reduce packing density (Renne and Ernst, 2023). Increasing the MUFA/SFA ratio therefore enhances membrane fluidity. In line with this, MUFA treatment or SCD1 overexpression reduced fluorescence polarization of the membrane dye TMA-DPH, indicating increased membrane fluidity. Thus, in Northeastern Asian brown frogs, cold-induced upregulation of SCD expression and enrichment of MUFA-containing phospholipids likely contribute to the maintenance of membrane structure and function under hypothermic conditions.

Reactive oxygen species are mediators of cold-induced injury (Magni et al., 1994; Rauen and de Groot, 1998). In addition to effects on membrane fluidity, we found that SFA treatment increased ROS production and lipid peroxidation, whereas MUFA treatment attenuated these responses under cold conditions. Oxidative stress and lipid peroxidation are key drivers of ferroptosis (Dixon and Olzmann, 2024). Previous studies have shown that exogenous MUFA supplementation suppresses lipid peroxidation and inhibits ferroptosis (Magtanong et al., 2019), and that SCD1 overexpression enhances resistance to ferroptosis, albeit with implications for tumorigenesis (Sen et al., 2023). In our study, the protective effect of MUFAs requires several hours of pre-treatment at 37□°C, suggesting that MUFAs act indirectly through cellular metabolic processes rather than exerting an immediate effect. The anti-ferroptotic action of MUFAs has been linked to acyl-CoA synthetase long-chain family member 3 (ACSL3), which facilitates incorporation of MUFAs into membrane phospholipids, thereby displacing peroxidation-prone polyunsaturated fatty acids (Magtanong et al., 2019). Notably, we also observed activation of necroptotic pathways during cold exposure, which was suppressed by MUFA treatment, indicating that MUFAs may preserve cellular viability by inhibiting both ferroptosis and necroptosis.

In summary, our findings demonstrate that MUFA supplementation enhances cell survival during low-temperature preservation of mouse liver and skin tissues, highlighting its translational potential for clinical tissue and organ preservation. Future studies are needed to evaluate the efficacy of MUFA-based interventions during rewarming and post-transplantation, and to further elucidate the molecular mechanisms by which MUFAs suppress lipid peroxidation and cell death pathways, including ferroptosis and necroptosis. Overall, this study identifies lipid monounsaturation as a key adaptive feature of cold tolerance and provides a comparative framework for leveraging evolutionary strategies to improve hypothermic tissue preservation.

## MATERIALS AND METHODS

### Animals

#### 1) Frog experiments

The African clawed frogs (*X. laevis*) used in this study were kindly provided by Professor Qinghua Tao (School of Life Sciences, Tsinghua University) and were 2-3-year-old males. Frogs were maintained in filtered, oxygenated purified water (pH 7.2) at 18-22 °C, with water changed every two days. Animals were fed an appropriate amount of bloodworms or beef liver every three days. Northeastern Asian brown frogs (*R. dybowskii*) were semi-artificially bred male frogs aged 2-3 years and were obtained from Jilin City, Jilin Province, China. These frogs were housed in food-grade plastic cylindrical containers (diameter, 57 cm; height, 75 cm) with clean stones placed at the bottom to provide shelter. Container tops were covered with sun-protective nets to minimize disturbance. Frogs were maintained at 18-22 °C in filtered, oxygenated purified water, with water replaced every two days, and were fed mealworms.

All animal procedures complied with ethical standards for scientific research, relevant regulations, and institutional guidelines. Prior to cold exposure, both *X. laevis* and *R. dybowskii* were acclimated at 20 °C for seven days to achieve metabolic equilibrium at ambient temperature. Frogs of each species were then randomly assigned to either a control group (20 °C) or a cold exposure group (1 °C), with 3-5 animals per group. Frogs in the cold exposure group were placed in ambient water and gradually cooled in a refrigerator to minimize acute cold shock. Control animals were maintained under normal feeding and water-changing schedules, whereas cold-exposed frogs were not fed but continued to receive regular water replacement. After the exposure period, all animals were euthanized by double pithing, and liver tissues were collected and stored at −80 °C for subsequent analyses.

#### 2) Mouse experiments

Wild-type C57BL/6J and ICR mice used in this study were bred in-house at the Tsinghua University Laboratory Animal Center. All animals were housed in a specific pathogen-free (SPF) facility, provided with a sterile pellet diet and water *ad libitum*, and maintained under a 12 h light/12 h dark cycle at 22-26 °C. All procedures were performed in accordance with an Institutional Animal Care and Use Committee (IACUC)-approved protocol (Approval No. 23-DHT24-1).

For time-course experiments assessing cold preservation-induced liver tissue degradation, 5-week-old female C57BL/6J mice were euthanized by cervical dislocation, followed by whole-body perfusion with ice-cold lactated Ringer’s solution. Intact livers were excised and placed in 10 mL ice-cold DMEM at 4 °C for the indicated preservation periods. For experiments involving palmitoleic acid (POA) treatment during liver and skin cold preservation, 4-week-old male ICR mice were injected via the tail vein with 250 µL of 5 mM POA (dissolved in DMEM after complexing with 10% BSA). Six hours after injection, mice were euthanized and subjected to whole-body perfusion with University of Wisconsin (UW) solution containing 250 µM POA. Livers were then excised and placed in 10 mL ice-cold UW solution supplemented with 250 µM POA at 4 °C.

### Proteomic analysis

#### 1) Sample preparation

Sample preparation for proteomics analysis was performed as previously described (Liu et al., 2024). Briefly, approximately 20 mg of liver tissue per sample was processed, with three biological replicates per group. Samples were lysed on ice in RIPA buffer supplemented with protease inhibitors. Ice-cold sonication was performed using a contact-type sonicator until the lysates became clear and non-viscous. Lysates were centrifuged at 12,000 rpm for 10 min at 4 °C, and the supernatants were collected. Five volumes of pre-chilled acetone were added, gently mixed, and incubated at −20 °C overnight to precipitate proteins. Precipitated proteins were pelleted by centrifugation at 7,500 rpm for 10 min, the supernatant was carefully removed, and residual acetone was allowed to air-dry. Protein pellets were dissolved in 300 µL of 8 M urea, and protein concentration was determined using a BCA assay.

For each sample, 100 µg of protein was adjusted to a final volume of 100 µL with 8 M urea. Under dark conditions, tris(2-carboxyethyl)phosphine (TCEP; final concentration 10 mM) and chloroacetamide (CAA; final concentration 40 mM) were added, mixed thoroughly, and incubated at room temperature for 60 min. The reaction was quenched by light exposure, after which 700 µL of PBS was slowly added to dilute the urea. Proteins were digested by adding MS-grade trypsin (2 µg; enzyme-to-substrate ratio 1:50, w/w) and incubating at 37 °C for 12 h. Peptide solutions were centrifuged at 12,000 rpm for 10 min, and the supernatants were collected. Trifluoroacetic acid (TFA) was added to a final concentration of 0.4% to terminate digestion and adjust the pH below 2.

Peptides were desalted using Waters Sep-Pak C18 solid-phase extraction (SPE) cartridges (50 mg capacity). Cartridges were sequentially activated with 1 mL of Solution 1 (acetonitrile), Solution 3 (0.5% acetic acid, 50% acetonitrile), and Solution 4 (0.1% TFA). Samples were loaded, washed with Solution 4 and Solution 2 (0.5% acetic acid), and eluted with 1.5 mL of Solution 3. Eluates were dried in a centrifugal evaporator and re-dissolved in 50 µL of 100 mM triethylammonium bicarbonate (TEAB).

TMT 6-plex reagents (5 mg) were dissolved in 250 µL of acetonitrile, and 15 µL of the reagent was added to each sample. Labeling reactions were incubated at room temperature for 60 min to ensure complete labeling. Following an additional 15 min incubation, all TMT-labeled samples were combined, acetonitrile was removed by centrifugal evaporation, and the total volume was adjusted to 1 mL with MS-grade water. After acidification to pH <2 with TFA, peptides were desalted using a single Sep-Pak C18 cartridge (100 mg capacity) and eluted with 1.5 mL of Solution 3. Eluates were concentrated to 100–200 µL by centrifugal evaporation and centrifuged at 12,000 rpm for 10 min. Supernatants were transferred to HPLC vials for fractionation.

High-pH reverse-phase fractionation was performed using an HPLC system equipped with a C18 column. Mobile phase A consisted of HPLC-grade water (pH 10 adjusted with ammonia), and mobile phase B consisted of 98% acetonitrile (pH 10 adjusted with ammonia). Peptides were separated using a 60 min linear gradient at 45 °C and monitored at 214 nm. Fractions were collected across the gradient, yielding 47 fractions, which were dried and concatenated into 12 final fractions. Dried peptides were re-dissolved for subsequent LC-MS/MS analysis.

#### 2) Mass spectrometry data acquisition and analysis

Peptide fractions were analyzed by liquid chromatography–tandem mass spectrometry (LC-MS/MS) using an EASY-nLC 1200 nanoLC system coupled to a Thermo Scientific Fusion Lumos Tribrid mass spectrometer. Peptides were separated on a C18 reverse-phase column using mobile phase A (0.1% formic acid in water) and mobile phase B (80% acetonitrile with 0.1% formic acid) at a flow rate of 300 nL/min. Peptides were ionized by electrospray ionization prior to mass spectrometric analysis.

Data were acquired in data-dependent acquisition (DDA) mode. Full MS1 scans were collected in the Orbitrap over a mass range of 350–1500 m/z at a resolution of 120,000 (at m/z 200), with an automatic gain control (AGC) target of 4 × 10□ and a maximum injection time of 50 ms. The top 40 most intense precursor ions from each MS1 scan were selected for MS2 analysis. Fragmentation was performed by higher-energy collisional dissociation (HCD), and fragment ions were detected in the Orbitrap at a resolution of 50,000 (at m/z 200), with an AGC target of 1 × 10□, a maximum injection time of 105 ms, and an isolation window of 1.6 Da. TMT 6-plex reporter ions (m/z 126–131) were detected in MS2 spectra for relative quantification. Raw data files were saved in *.raw format for downstream analysis.

Raw MS data were processed for peptide and protein identification, quantification, and post-translational modification analysis using Proteome Discoverer (version 3.2) with the Sequest HT search engine. Species-specific UniProt databases were used: Xenopus laevis (African clawed frog; TaxID 8355) for African clawed frog samples and Rana temporaria (Common frog; TaxID 8407) for Northeast Asian brown frog samples. Databases were downloaded as FASTA files and imported into the software. Trypsin was specified as the proteolytic enzyme, allowing up to two missed cleavages and a minimum peptide length of six amino acids. Fixed modifications included TMT 6-plex labeling on peptide N-termini and lysine residues, as well as carbamidomethylation of cysteine. Variable modifications included methionine oxidation and protein N-terminal acetylation. Precursor ion mass tolerance was set to 10 ppm, and fragment ion tolerance was set to 0.6 Da. Protein quantification was based on TMT reporter ion intensities, and data were normalized to total peptide abundance. For acetylation analyses, identical DDA parameters were applied, and relative peptide abundances were quantified based on MS1 peak areas. The visualization of the proteomics analysis results was performed using SRplots (Tang et al., 2023) platform.

### Metabolomic analysis

#### 1) Sample preparation

Sample preparation and data acquisition for untargeted metabolomic analysis was performed as previously described (Tang et al., 2016). Briefly, approximately 90 mg of liver tissue per sample was homogenized in1 mL 80% (v/v) HPLC-grade methanol. The homogenates were then shaken for 1 min at 4 °C and incubated at −80 °C for 2 h. Samples were subsequently centrifuged at 14,000 × g for 20 min at 4 °C. Equal volumes of the supernatants were transferred to new microcentrifuge tubes and dried to a pellet using a vacuum concentrator (SpeedVac). Dried samples were stored at −80 °C until metabolomic analysis.

#### 2) LC-MS/MS data acquisition and analysis

For positive ion mode analysis, chromatographic separation was performed on a BEH amide column (2.1 × 100 mm, 1.7 µm; Waters) at 35 °C. Mobile phase A consisted of 0.63 g ammonium formate dissolved in 50 mL HPLC-grade water, followed by the addition of 950 mL HPLC-grade acetonitrile and 1 µL formic acid. Mobile phase B consisted of 0.63 g ammonium formate dissolved in 500 mL HPLC-grade water, followed by the addition of 500 mL HPLC-grade acetonitrile and 1 mL formic acid. The initial mobile phase composition was 99% A and 1% B, with a flow rate of 300 µL/min.

For negative ion mode analysis, a BEH C18 column (2.1 × 100 mm, 1.7 µm; Waters) was used at 40 °C. Mobile phase A consisted of 0.3953 g ammonium bicarbonate dissolved in 1 L HPLC-grade water (pH ∼8), and mobile phase B was 100% acetonitrile. The initial mobile phase composition was 99% A and 1% B, with a flow rate of 200 µL/min. Untargeted metabolomic profiling was performed on a Q Exactive HF Orbitrap mass spectrometer (Thermo Scientific), calibrated according to the manufacturer’s instructions.

Metabolites were identified using TraceFinder 3.2 (Thermo Scientific) in combination with an internal database of more than 1,500 endogenous metabolites curated by the Metabolomics and Lipidomics Core Facility at Tsinghua University. Mass tolerances were set to 10 ppm for precursor ions and 15 ppm for fragment ions. Metabolite identification was achieved at two confidence levels: one confirmed by MS/MS spectral matching and one based solely on accurate precursor mass. Relative quantification was performed using chromatographic peak areas, allowing a retention time shift of up to 0.25 min for peak alignment. The visualization of the metabolomics analysis results was performed using SRplots (Tang et al., 2023) and MetaboAnalyst 6.0 (Pang et al., 2024) platforms.

### Lipidomic analysis

#### 1) Sample preparation

Sample preparation and data analysis for untargeted lipidomics was performed as previously described (Xu et al., 2018). Briefly, approximately 50 mg of tissue from the same region of each liver was homogenized in 600 µL methanol. Following homogenization, 2 mL of methyl tert-butyl ether (MTBE) was added, and samples were vortexed thoroughly. The suspension was divided into two equal aliquots (1,200 µL each) and transferred into two 1.5 mL microcentrifuge tubes. Subsequently, 250 µL of water was added to each aliquot to induce phase separation. Vortexing and settling were repeated three times. After extraction, samples were centrifuged at 8,000 rpm for 15 min at room temperature. An equal volume (960 µL) of the upper organic phase was transferred into new 1.5 mL tubes. Organic solvents were evaporated under a gentle stream of nitrogen gas without heating until a transparent lipid film remained. Dried samples were stored at −80 °C prior to analysis.

#### 2) LC-MS/MS data acquisition and analysis

Reversed-phase liquid chromatography was performed using a CORTECS C18 column (2.1 × 100 mm, 2.7 µm; Waters). Mobile phase A consisted of 400 mL HPLC-grade water containing 0.77 g ammonium acetate mixed with 600 mL HPLC-grade acetonitrile, and mobile phase B consisted of 10% acetonitrile and 90% isopropanol (v/v). Separation was carried out at a flow rate of 0.25 mL/min with the column temperature maintained at 40 °C. Data acquisition was performed on an Orbitrap Exploris 240 mass spectrometer (Thermo Fisher Scientific) coupled to an ultra-high-performance liquid chromatography (UHPLC) system (Vanquish, Thermo Fisher Scientific).

Lipid identification was conducted using LipidSearch 4.2 (Thermo Fisher Scientific), which contains a database comprising 18 lipid classes and more than 1.5 million fragment ions. Lipids were identified based on MS/MS spectral matching. Mass tolerances were set to 8 ppm for precursor ions and 15 ppm for fragment ions. The m-score threshold was set to 5, and lipids were filtered according to quality scores (A, B, C, and D). All 71 lipid subclasses in the database were included for identification. In positive ion mode, +H and +NH₄⁺ adducts were considered, whereas in negative ion mode, −H and +CH₃COO⁻ adducts were used due to the presence of ammonium acetate in the mobile phase. Only lipids with chromatographic peak areas greater than 5 × 10□ were retained for reliable identification. Quantification allowed a retention time tolerance of 0.25 min.

### Cell culture and treatments

293T, BV2, and HeLa cells were cultured in DMEM, whereas K562 cells were cultured in RPMI 1640 medium; all media were supplemented with 10% fetal bovine serum (FBS) and 1% penicillin–streptomycin. Cells were maintained at 37 °C in a humidified incubator with 5% CO₂, and routine mycoplasma testing was performed to ensure cultures were contamination-free.

Fatty acid stock solutions were prepared by dissolving fatty acids in 100% ethanol, vortexing until fully dissolved, adjusting to a final concentration of 250 mM, and storing at −20 °C. To prepare fatty acid working solutions, a 10% (w/w) BSA solution was first prepared by dissolving BSA in the corresponding culture medium. The 250 mM fatty acid stock solution was then diluted 1:100 into the 10% BSA solution to yield a final concentration of 2.5 mM. The mixture was incubated at 37 °C for 30 min with occasional vortexing to ensure complete complex formation.

For fatty acid treatment, the working solution was added to fresh complete culture medium 6 h prior to cold exposure. Cells were incubated in fatty acid-supplemented medium at 37 °C for 6 h and subsequently transferred to a 4 °C refrigerator for cold exposure.

### Construction of gene overexpression cell lines

Mouse CRISPR activation (mCRISPRa) lentiviral plasmids, including lenti-EF1a-dCas9-VP64 (blasticidin-resistant), lenti-EF1a-MCP-p65-HSF1 (hygromycin B-resistant), and pklv-mU6-3 ′ 2ms2-pgk-puro-2a-bfp (newSAMem; puromycin-resistant), were kindly provided by Professor Yu Zhang (Chinese Institutes for Medical Research, Beijing). Lentiviruses expressing dCas9-VP64 and MCP-p65-HSF1 were packaged in 293T cells using a second-generation lentiviral system (psPAX2 and pMD2.G) with Lipofectamine 3000 (Invitrogen, Cat# L3000001). Viral supernatants were collected and used to sequentially infect BV2 cells. Infected cells were selected with blasticidin (5 mg/L; Cat# ST018, Beyotime) and hygromycin B (50 mg/L; Cat# ST1389, Beyotime) to generate a BV2 CRISPRa cell line stably expressing both dCas9-VP64 and MCP-p65-HSF1.

Target sequences for the Scd1 promoter were cloned into the mCRISPRa library plasmid pklv-mU6-3’2ms2-pgk-puro-2a-bfp (newSAMem) using Gibson assembly. Three high-scoring Scd1-targeting single-guide RNA (sgRNA) sequences from the Jonathan Weissman laboratory mCRISPRa-v2 library (https://weissman.wi.mit.edu/crispr/) and one non-targeting control were selected: sgScd1#1 (gaagccaggcgtggaggtga), sgScd1#2 (ggatgctgaaggcatcaaca), sgScd1#3 (gagagggcgggaccaagaac), and sgNC (gcggtgtggcgtcctgtatt). Complementary DNA oligonucleotides were annealed to generate double-stranded inserts and ligated into the linearized newSAMem plasmid by Gibson assembly. Correct plasmid construction was verified by Sanger sequencing.

Lentiviruses carrying sgRNA constructs were subsequently packaged and used to infect BV2 CRISPRa cells, followed by puromycin selection (1 mg/L; Cat# ST551, Beyotime) to establish stable Scd1-overexpressing cell lines.

### Cell viability assay

Cell viability and death were assessed using Calcein AM/propidium iodide (PI) double staining (Cat# C2015, Beyotime). Culture medium was gently aspirated, and cells were washed once with PBS. Calcein AM/PI staining solution was prepared according to the manufacturer’s instructions using phenol red-free DMEM. A total of 100 µL of staining solution was added to each well of a 96-well plate, followed by incubation at 37 °C for 30 min in the dark. Fluorescence imaging was then performed (Calcein AM: λ_ex/λ_em = 494/517 nm; PI–DNA complex: λ_ex/λ_em = 535/617 nm), and the ratio of live (Calcein AM-positive) to dead (PI-positive) cells was calculated.

### ROS measurement

Intracellular reactive oxygen species (ROS) levels were measured using DCFH-DA (Cat# G1706, Servicebio). Cells were collected by centrifugation at 1,000 rpm at 4 °C for 3 min, washed once with PBS, and resuspended in DCFH-DA working solution at a density of 1–5 × 10□ cells/mL. Cells were incubated at 37 °C in the dark for 30 min with gentle shaking every 5 min. After incubation, cells were washed twice with PBS and analyzed by flow cytometry (λ_ex/λ_em = 488/525 nm).

### Lipid peroxidation measurement

Lipid peroxidation was assessed using BODIPY 581/591 C11 (Cat# S0043, Beyotime). For microscopy, culture medium was removed and cells were washed once with PBS. For flow cytometry, cells were collected by centrifugation at 1,000 rpm at 4 °C for 3 min, washed once with PBS, and resuspended in 2 µM BODIPY 581/591 C11 working solution. Cells were incubated at 37 °C in the dark for 30 min with gentle shaking every 5 min. Following incubation, cells were washed twice with PBS. Lipid peroxidation levels were analyzed by confocal microscopy or flow cytometry, with the reduced form detected at λ_ex/λ_em = 581/591 nm and the oxidized form at λ_ex/λ_em = 488/510 nm.

### MDA measurement

Malondialdehyde (MDA) levels were measured using a commercial assay kit (Cat# BC0025, Solarbio). Cells were lysed in 1 mL of extraction buffer per 5 × 10□ cells (or per 100 mg of tissue), sonicated, and centrifuged at 8,000 × g at 4 °C for 10 min. Supernatants were collected on ice. Detection reagents were added according to the manufacturer’s instructions, followed by incubation at 95 °C for 1 h. Samples were then cooled on ice, centrifuged at 10,000 × g for 10 min at room temperature, and 200 µL of the supernatant was transferred to a 96-well plate for absorbance measurement at 532 nm and 600 nm.

### Western blotting

Cells were washed twice with ice-cold PBS, aspirated, and lysed on ice for 10 min in ice-cold RIPA buffer supplemented with a protease inhibitor cocktail (Selleck, Cat# B14001) and a phosphatase inhibitor cocktail (Selleck, Cat# B15001). Cells were collected using a cell scraper and sonicated. Lysates were centrifuged at 12,000 rpm at 4 °C for 10 min, and the supernatants were transferred to new tubes. Protein concentrations were determined using a BCA assay (Cat# P0009, Beyotime) and equalized with complete RIPA buffer. LDS sample buffer containing 5% β-mercaptoethanol (Invitrogen, Cat# NP0007) was added, and samples were heated at 70 °C for 10 min to denature proteins. Samples were either used immediately for SDS-PAGE or stored at −20 °C.

SDS-PAGE gels were prepared according to the molecular weights of target proteins. Proteins were transferred to PVDF membranes, blocked with skim milk for 1 h, and incubated with primary antibodies overnight at 4 °C. Membranes were then incubated with appropriate secondary antibodies for 2 h at low speed. Chemiluminescent signals were detected using a ChemiDoc imaging system (Bio-Rad).

### Measurement of cell membrane fluidity

Cell membrane fluidity was measured using the fluorescent probe TMA-DPH (Ex/Em = 355/430 nm; Maclin, Cat# T911697). Cells were collected by centrifugation at 1,000 rpm at 4 °C for 3 min, washed once with PBS, and resuspended in 1 µM TMA-DPH working solution. Cells were incubated at 37 °C in the dark for 15 min, washed twice with PBS, and resuspended in PBS. Fluorescence polarization was measured using a Revvity EnVision I plate reader (λ_ex/λ_em = 355/460 nm).

### H&E and TUNEL staining

Tissues were fixed in 4% paraformaldehyde (PFA) at 4 °C for 24 h and subsequently embedded in paraffin. Sections (5 µm thick) were cut using a microtome, floated on a 40 °C water bath, mounted onto anti-drop slides, and deparaffinized. Hematoxylin and eosin (H&E) staining was performed according to standard procedures, followed by coverslipping. Whole-slide images were acquired using a 3DHISTECH Pannoramic SCAN scanner at 20× magnification.

For TUNEL staining, paraffin-embedded sections were deparaffinized and rehydrated. After gentle drying, a hydrophobic barrier was drawn around the tissue using a histology pen. Proteinase K working solution was applied to cover the tissue, and sections were incubated at 37 °C for 22 min for antigen retrieval. Sections were washed three times with PBS, gently dried, and permeabilized with 0.1% Triton X-100 in PBS at room temperature for 20 min. The TUNEL reaction mixture was prepared using a commercial kit (Servicebio, Cat# G1502) by mixing terminal deoxynucleotidyl transferase (TdT) enzyme, dUTP, and buffer at a 1:5:50 ratio. The reaction mixture was applied within the hydrophobic barrier, and sections were incubated in a humidified chamber at 37 °C for 2 h. Sections were then washed with PBS and counterstained with DAPI for 10 min. Coverslips were mounted using anti-fade mounting medium. Whole-slide fluorescence images were acquired using an Axio Scan.Z1 microscope (Zeiss) at 20× magnification (DAPI: λ_ex/λ_em = 340/488 nm; Cy3: λ_ex/λ_em = 554/568 nm).

### Measurement of transaminase levels

Alanine aminotransferase (ALT) and aspartate aminotransferase (AST) levels released into University of Wisconsin (UW) solution from liver tissues were measured using commercial ALT and AST assay kits (Sjodax, Beijing) with an automated biochemical analyzer (Hitachi 7100).

### Statistical analyses

All experiments were performed with at least three independent biological replicates unless otherwise indicated. Data are presented as mean ± standard deviation (SD) or mean with 95% confidence interval (CI), as specified in the figure legends. Statistical analyses were conducted using GraphPad Prism 8 (GraphPad Software, San Diego, CA, USA). Kaplan-Meier survival analysis was used to compare survival curves. Comparisons between two groups were performed using an unpaired two-tailed Student’s t-test. For comparisons involving three or more groups, two-way ANOVA was applied, followed by Sidak’s post hoc multiple-comparison test. A p value < 0.05 was considered statistically significant.

### Data availability

Proteomics data have been deposited to the ProteomeXchange Consortium via the iProX partner repository with the dataset identifier PXD073918. Metabolomics data have been deposited in the MetaboLights repository with the accession No. MTBLS13816 (Reviewer link: https://www.ebi.ac.uk/metabolights/reviewerf62530b4-59bf-48c4-a6a9-8f5aa902f7ad).

## ACKNOWLEDGMENTS

We thank Professor Yu Zhang and Xiaofang Si from Capital Medical University for providing CRISPRa plasmids; Dr. Ying Li from the Tsinghua University Cryo-EM facility for assistance with experiments; Professor Jie Yan and Jinze Yang from Peking University Third Hospital for discussions; and the Tsinghua University Laboratory Animal Center for supporting the mouse experiments. This study was supported by the National Natural Science Foundation of China (Grant Nos. T2293763 and 82301607), the Beijing Natural Science Foundation (Grant No. IS24040), the Open Research Fund of the State Key Laboratory of Complex, Severe, and Rare Diseases (Grant No. 2025-I-PY-001), the China Postdoctoral Science Foundation (Grant No. 2024M751632), and Tsinghua University Shuimu Scholar Program.

## AUTHOR CONTRIBUTIONS

H.D. and R.Z. conceptualized and directed the study. R.Z., W.H., and H.D. designed and performed the experiments. Y.C., T.N., Y.T., Z.L., C.Z., Y.J., X.L., Y.L., J.L., Z.L., J.W., and Q.T. contributed to the execution, support, and analysis of experiments, data interpretation and/or advice, and wrote the paper.

## DECLARATION OF INTERESTS

The authors declare no competing financial interests except that a patent application (No. 202510320558.6) related to this work has been filed in China.

## Notes

https://proteomecentral.proteomexchange.org/ui?pxid=PXD073918

https://www.ebi.ac.uk/metabolights/reviewerf62530b4-59bf-48c4-a6a9-8f5aa902f7ad

### Competing Interest Statement

The authors have declared no competing interest.

