## Supplemental Figures S1-S7 for "Lipid monounsaturation confers cold tolerance and improves tissue viability during hypothermic storage"

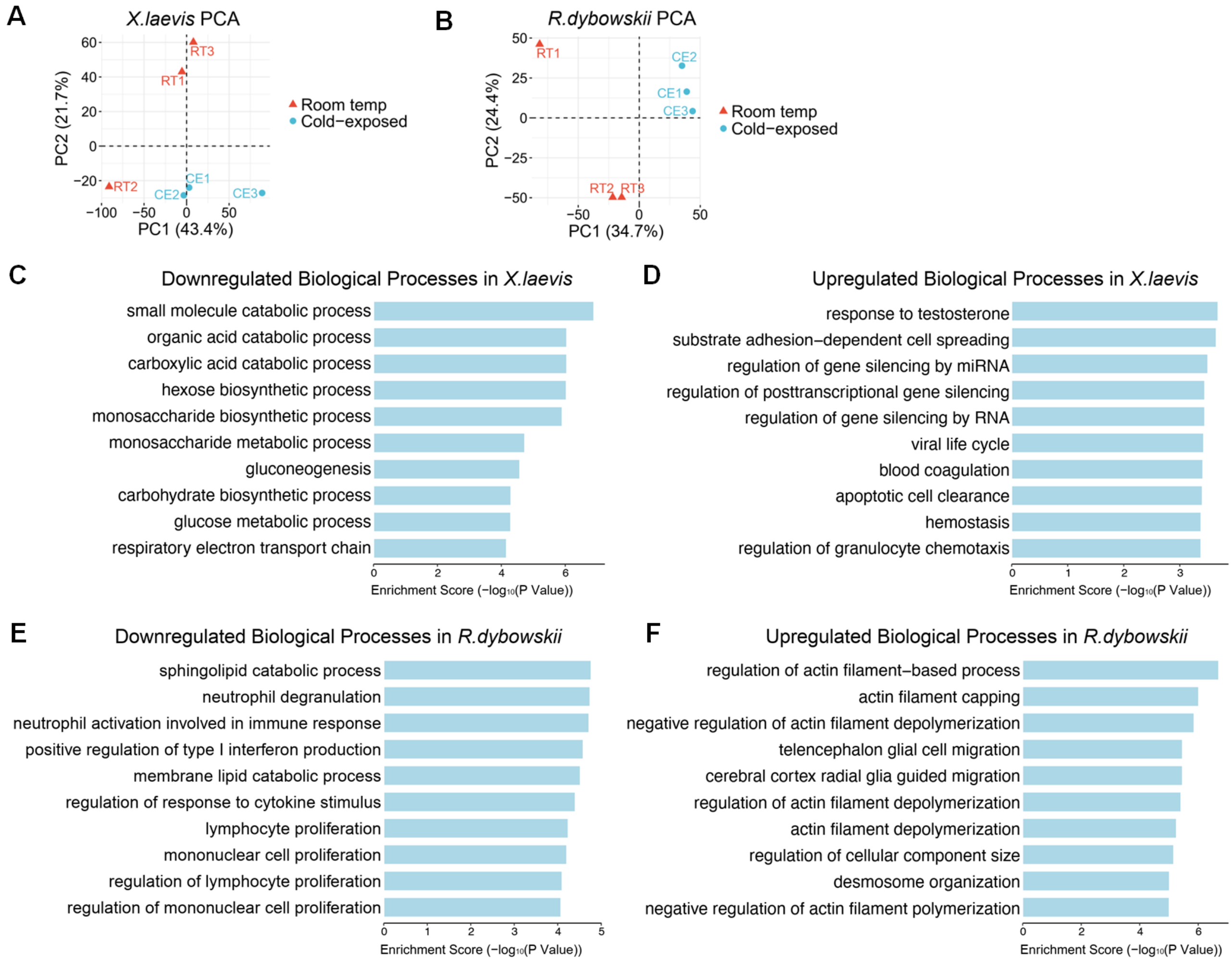


**Figure S1. Proteomic analysis of livers from two frog species upon cold exposure.**
(A and B) PCA of African clawed frog (A) and Northeastern Asian brown frog livers (B) showing the relative distances between samples.
(C and D) GO analysis showing the top 10 significantly enriched biological processes among upregulated (C) and downregulated (D) proteins in African clawed frog liver upon cold exposure.
(E and F) GO analysis showing the top 10 significantly enriched biological processes among upregulated (E) and downregulated (E) proteins in Northeastern Asian brown frog liver upon cold exposure.

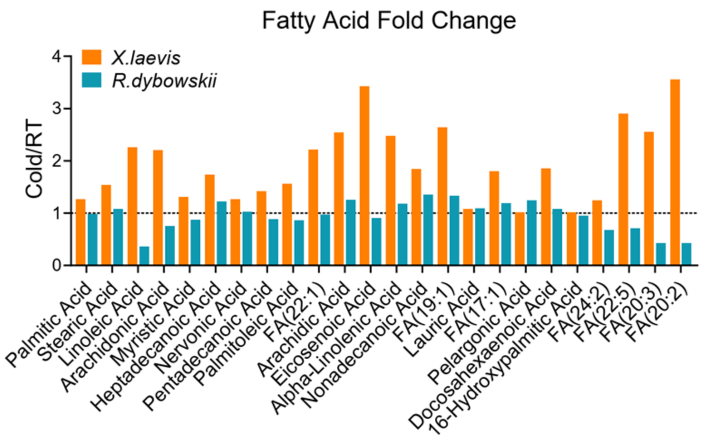


**Figure S2. Changes in free fatty acids in the livers from two frog species upon cold exposure.** Fold changes of free fatty acids after cold exposure in African clawed frog (*X. laevis*, orange bars) and Northeastern Asian brown frog (*R. dybowskii*, blue bars) liver.

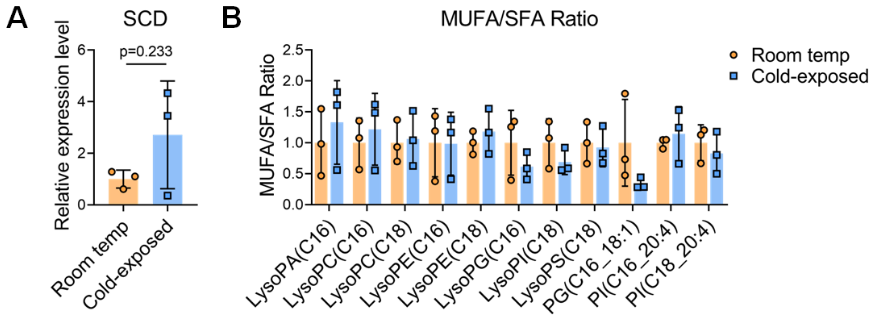


**Figure S3. SCD expression and phospholipid monounsaturation in African clawed frog liver.**
(A) Relative SCD protein expression levels in African clawed frog liver measured by quantitative proteomic analysis. N = 3.
(B) Relative ratios of monounsaturated to saturated phospholipids (MUFA/SFA) in African clawed frog liver quantified through metabolomic analysis. N = 3.
Data are presented as mean ± SD. Statistical significance was determined using unpaired Student's t-test.

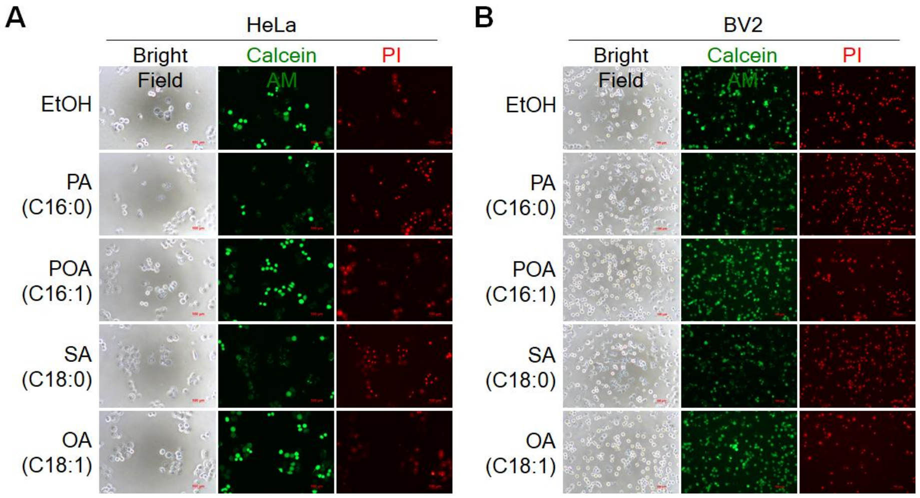


**Figure S4. MUFA improves cell survival under cold conditions.**
Calcein AM/PI double staining and representative microscopy imaging showing survival (green) and death (red) of HeLa (A) and BV2 (B) cells pretreated with different fatty acids followed by cold exposure.

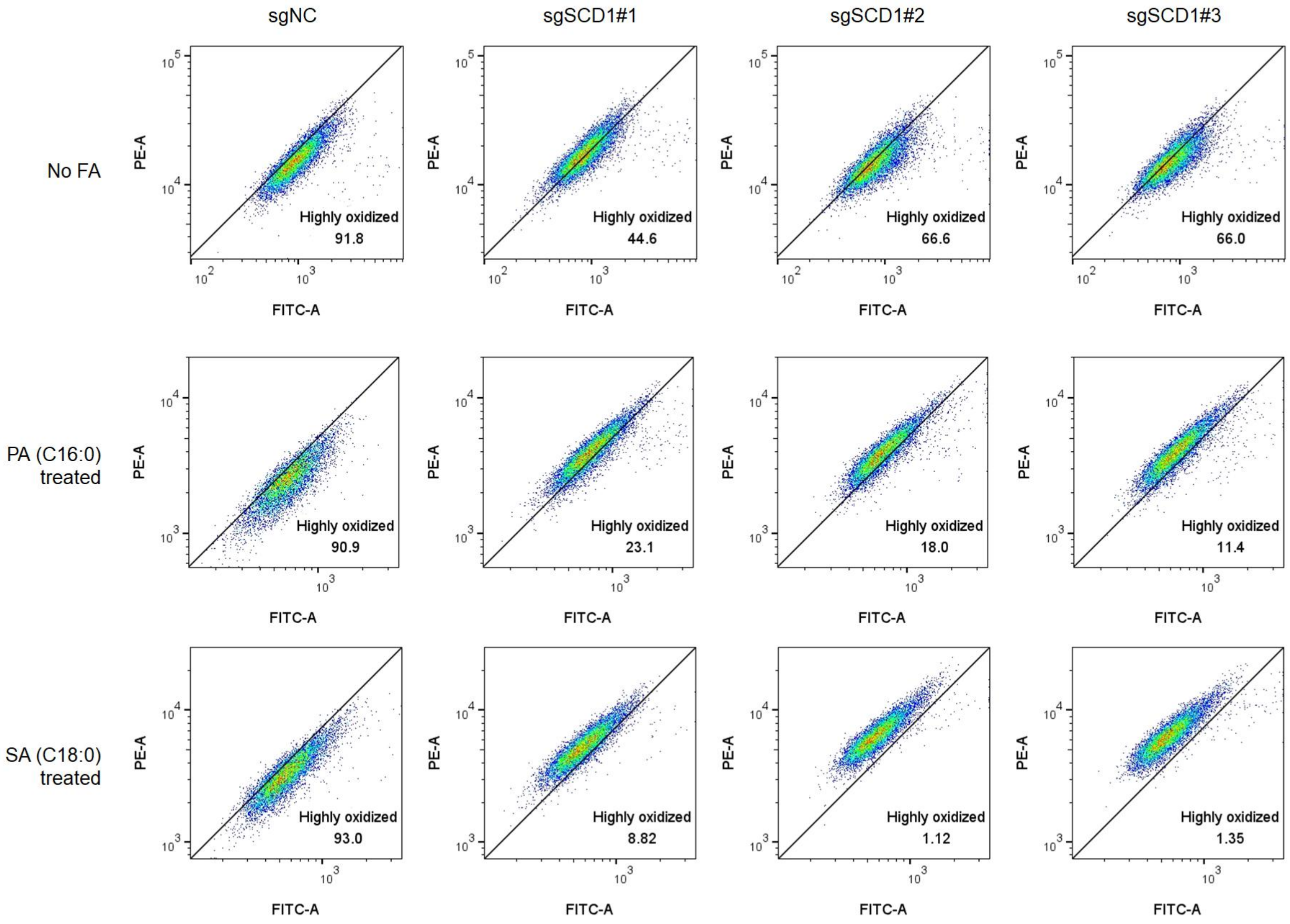


**Figure S5. Effects of SCD1 overexpression on lipid peroxidation in BV2 cells under cold exposure.**

Flow cytometry of lipid peroxidation after cold exposure with or without PA or SA pretreatment. X-axis (FITC): oxidized BODIPY; Y-axis (PE): reduced BODIPY. Bottom right shows the proportion of highly oxidized cells.

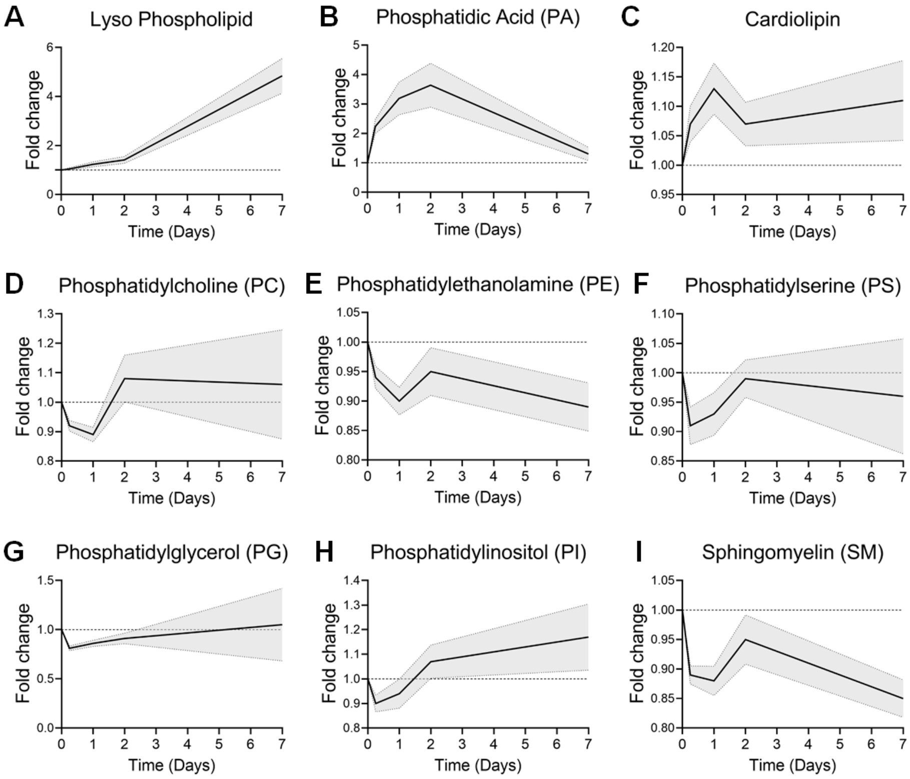


**Figure S6. Lipidomic analysis of mouse liver during cold storage.**
Levels of lysophospholipids (A), phosphatidic acid (B), cardiolipin (C), phosphatidylcholine (D), phosphatidylethanolamine (E), phosphatidylserine (F), phosphatidylglycerol (G), phosphatidylinositol (H), and sphingomyelin (I) in mouse liver tissues over time during cold storage. Data in each panel are normalized to the total signal values and presented as mean with 95% CI.

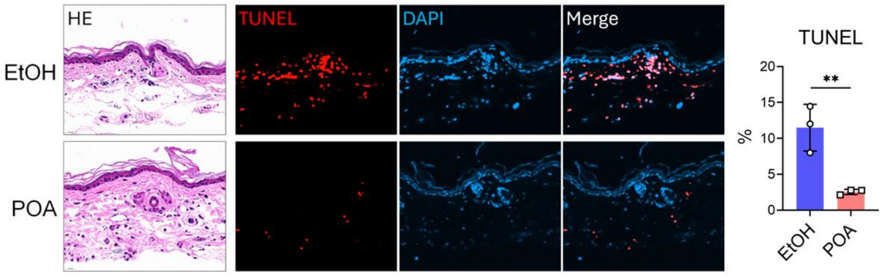


**Figure S7. POA treatment reduces apoptosis during mouse skin cold storage.**
HE and TUNEL imaging of mouse ear skin after 24 hours of cold storage treated with POA or ethanol vehicle. Nuclei were shown with DAPI staining. The percentages of TUNEL-positive cells were quantified. N = 3.
Data are presented as mean ± SD. Statistical significance was determined using unpaired Student's t-test, $^{**}p < 0.01$.
